# Gait speed and surface stiffness interactively modulate selected muscle- but not joint-synergy recruitment during compliant-surface walking

**DOI:** 10.64898/2026.05.13.725018

**Authors:** Toshihiro Ishii, Ken Takiyama

**Author notes:** Corresponding author: Ken Takiyama.

## Abstract

Humans adapt walking to different speeds and compliant surfaces, but whether gait speed and surface stiffness shape joint kinematics and muscle activity independently or interactively remains unclear. We reanalyzed an open variable-stiffness treadmill dataset collected at three speeds and four stiffness levels. Tensor decomposition extracted joint and muscle synergies and condition-specific recruitment coefficients. Stride length and stride time were modulated by speed and stiffness without interactions. Joint-synergy recruitment showed speed effects in all three components and stiffness effects in two components, but no speed-by-stiffness interactions. Among five muscle synergies, three were modulated without interaction, whereas two related to weight acceptance and forward propulsion showed significant speed-by-stiffness interactions. Their recruitment increased with speed, but stiffness-dependent differences decreased at higher speeds. These findings suggest that speed and stiffness largely modulate stride- and joint-level control independently, while interactively shaping selected muscle-synergy recruitment.

## INTRODUCTION

How the motor system controls the body remains a fundamental question in motor science. The musculoskeletal system has more degrees of freedom (DoFs) than are strictly necessary for many movement repertoires. This redundancy allows multiple patterns of muscle activity and joint motion to produce the same movement, requiring the motor system to select appropriate coordination patterns. One hypothesis for addressing this redundancy is the synergy framework^1–3^. In this framework, the motor system may reduce the effective DoFs by controlling groups of muscles and joints rather than each muscle or joint independently^3–7^. Here, groups of muscles are referred to as muscle synergies, their associated temporal components as activity profiles, groups of joints as joint synergies, and their associated temporal patterns as angular variations.

Muscle and joint synergies have been examined in various movements, including gait^5,8–10^, postural control^7,11,12^, and arm reaching^13^. The present study focuses on human gait, a representative whole-body dynamic movement. During gait at different speeds on hard, flat surfaces, 4-5 muscle synergies can explain approximately 90% of the variance in muscle activity^5,8,9^ and are consistent across a wide range of gait velocities^5,10^. Similarly, two joint synergies can be extracted from three lower-limb joint angles^3,9^ and remain consistent during walking and obstacle crossing^9^.

In addition to their consistency, the speed-dependent modulation of muscle and joint synergies has been investigated. Muscle activity, joint motion, and center-of-mass (CoM) trajectories change with walking speed^3,14,15^. Speed-dependent and mode-dependent modulations (i.e., walking versus running) have also been examined for muscle synergies^10,16–18^ and joint synergies^18^. Together, these studies characterize how muscle and joint synergies are organized during human gait on hard, flat surfaces.

In daily life, however, gait is adapted not only to speed but also to surface conditions, such as paved roads, ice, snow, and sand. Despite this ecological relevance, relatively few studies have examined synergy structure during gait on compliant surfaces. Previous work has investigated various aspects of compliant-surface walking, including CoM trajectories, muscle activity, and toe trajectories related to gait stability^19^; modulation of CoM trajectories and lower-limb dynamics by surface stiffness^20^; age-related differences in muscle synergies^21^; inter-leg coordination controlled by split-belt treadmill stiffness^22^; and predictive control on flexible surfaces^23^.

However, it remains unclear how muscle and joint synergies are modulated when both gait speed and surface stiffness vary, and specifically whether these factors affect synergies independently or interactively. Addressing this question requires gait speed to be controlled across all surface-stiffness conditions. Walking speed is a strong confounding factor in gait analysis, as demonstrated in comparisons between patients with knee osteoarthritis and healthy controls^24–26^. In previous studies of gait on compliant surfaces^19–21^, gait speed was not necessarily maintained at a constant level because walking was often examined during stepping onto mattresses or specialized surfaces with one or both legs. Consequently, the influence of gait speed could not be fully separated from the influence of surface compliance. The recent development of the Variable Stiffness Treadmill 2 (VST 2)^27^ enables measurements of muscle activity and joint angles across different surface stiffness levels while maintaining a constant gait speed, and vice versa. The open dataset collected with the VST 2 therefore provides an opportunity to test whether gait speed and surface stiffness modulate muscle and joint synergies independently or interactively.

A key methodological issue is how to analyze muscle activity and joint-angle data with multiple experimental factors. Muscle and joint synergies have typically been estimated using non-negative matrix factorization (NNMF) and principal component analysis (PCA), respectively^3,5,7,10,28^. These matrix factorization methods are appropriate for two-factor representations, such as spatial and temporal components: muscle synergies and activity profiles, or joint synergies and angular variations. However, to evaluate the effects of gait speed and surface stiffness, the analysis must account for more than two factors: spatial components, temporal components, and condition-dependent modulations. We therefore used tensor decomposition^29^, which is designed for data with more than two factors. Specifically, we used CANDECOMP/PARAFAC (CP) decomposition because it is a straightforward extension of NNMF and PCA from the perspective of interpretation: it estimates synergies, their temporal components, and their recruitment in each experimental condition. We have previously applied CP decomposition to motion data^30,31^, including gait, to extract task dependency and individual differences. This approach revealed speed-cand mode-dependent modulations of muscle and joint synergies^18^, a muscle synergy specific to gait in people with knee osteoarthritis as speed increased^26^, and adaptation-dependent modulation of muscle synergies during split-belt treadmill adaptation^32^. These findings support the use of CP decomposition to examine speed- and surface-stiffness-dependent modulation of muscle and joint synergies.

In the present study, we reanalyzed a recent open dataset^33^ that includes joint-angle and electromyography (EMG) data collected during VST 2 walking at controlled gait speeds and surface stiffness levels. CP decomposition allowed us to extract speed- and surface-stiffness-dependent modulations as scalar recruitment coefficients. For example, if a synergy is not recruited at a given gait speed in a given participant, the associated coefficient is expected to be close to zero; if the recruitment of that synergy increases in proportion to gait speed, the coefficients for each speed are expected to increase accordingly^18^. We applied repeated-measures ANOVA (rANOVA) to these recruitment coefficients, with gait speed and surface stiffness as within-subject factors, to test whether these factors affect muscle and joint synergies independently or interactively.

We first analyzed stride length and stride time across gait speeds and surface stiffness levels. We then extracted joint and muscle synergies using CP decomposition. Finally, we applied rANOVA to the recruitment coefficients of each synergy to evaluate the effects of gait speed, surface stiffness, and their interaction.

## RESULTS

### Controlled-speed compliant-surface walking data were prepared for tensor decomposition

We reanalyzed an open dataset^33^ in which electromyography (EMG) and joint-angle data were recorded during treadmill walking under 12 conditions: four surface-stiffness levels (25, 40, 80, and 1000 kN/m) at three walking speeds (0.6, 0.8, and 1.0 m/s; Figure 1A). For each condition, we analyzed steady-state gait by selecting the last 30 gait cycles that satisfied the inclusion criteria from the latter 100 gait cycles (Figure 1A, see Methods for the inclusion criteria). Stride parameters, joint angles, and EMG profiles were averaged across these gait cycles in each participant and condition before tensorization (Figures 1B and 1C). Six participants were excluded because their EMG signals did not satisfy the quality-control criteria (Figure S1); the final analyses therefore included 14 participants (7 female and 7 male; age range, 19-37 years).

**Figure 1.**
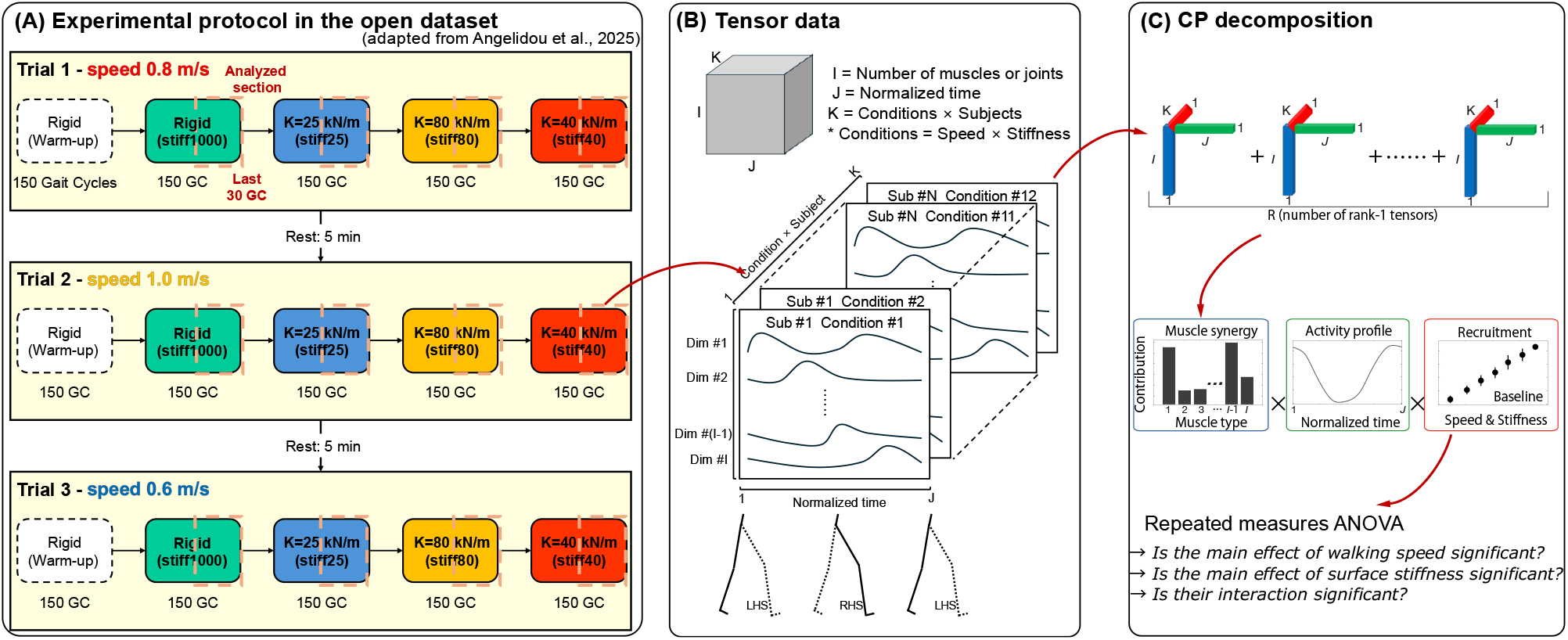
Analysis workflow for tensor-decomposition-based synergy analysis. (A) Experimental protocol in the open dataset, adapted from Angelidou et al.^33^. Participants walked at three speeds (0.6, 0.8, and 1.0 m/s) and four surface-stiffness levels (25, 40, 80, and 1000 kN/m); the present analysis used the last 30 qualifying gait cycles from each condition. (B) EMG and joint-angle data were averaged across the selected gait cycles. Before tensorization, joint-angle data were standardized within each participant and joint across all conditions, and EMG data were min-max normalized within each participant and muscle across all conditions. The EMG tensor was 10 x 200 x 168 and the joint-angle tensor was 6 x 100 x 168, where the first dimension denotes muscles or joints, the second denotes normalized time points, and the third denotes 14 participants x 12 conditions. (C) CP decomposition was applied separately to the joint-angle and EMG tensors to extract spatial synergies, temporal components, and condition-specific recruitment coefficients. Repeated-measures ANOVA was then used to test the main effects of walking speed and surface stiffness and their interaction.

### Stride length and stride time were independently modulated by speed and stiffness

We first examined whether walking speed and surface stiffness interactively modulated stride parameters (Figure 2). Stride length increased with walking speed (F(2, 26) = 414.0, p = 1.9× 10^−20^) and with decreasing surface stiffness (F(3, 39) = 6.2, p = 0.0014). The speed-by-stiffness interaction was not significant (F(6, 78) = 1.2, p = 0.32). Stride time decreased as walking speed increased (F(2, 26) = 157.3, p = 3.0 × 10^−15^) and increased on softer surfaces (F(3, 39) = 4.9, p = 0.0053), again without a significant interaction (F(6, 78) = 0.85, p = 0.53). These results indicate that walking speed and surface stiffness affected stride parameters independently rather than interactively.

**Figure 2.**
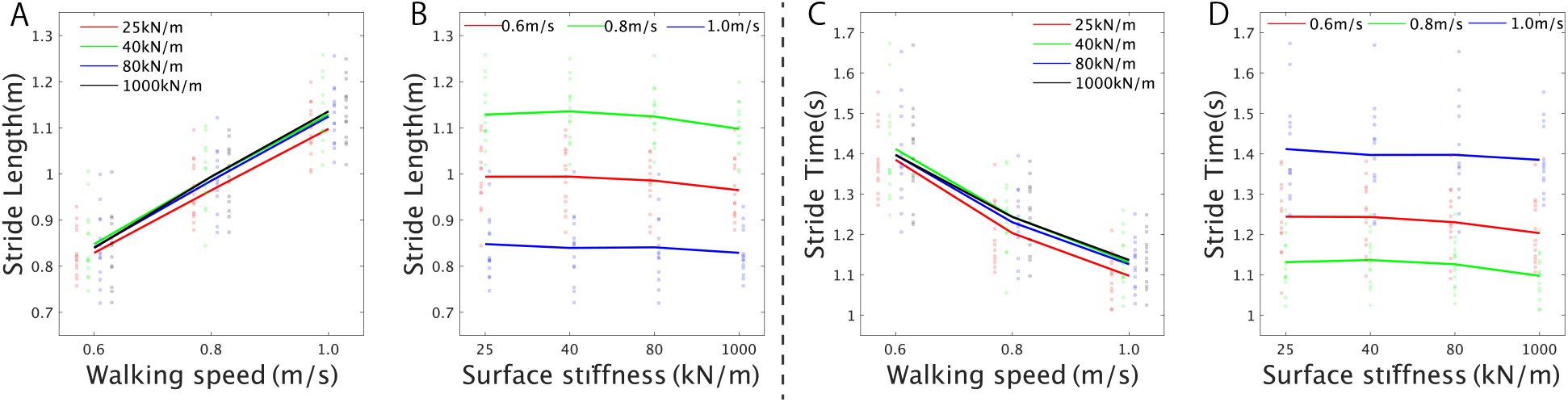
Stride parameters are modulated independently by walking speed and surface stiffness. (A and B) Stride length plotted by walking speed (A) and surface stiffness (B). (C and D) Stride time plotted by walking speed (C) and surface stiffness (D). Colored lines show means across participants, and transparent points show individual participant-condition values. In (A and C), colors denote surface stiffness; in (B and D), colors denote walking speed. Two-way repeated-measures ANOVA (n = 14) showed significant main effects of walking speed and surface stiffness for both stride length and stride time, with no significant speed-by-stiffness interaction. For stride length: walking speed, F(2,26)= 414.0, *p* = 1.9× 10^− 20^; surface stiffness, F(3,39)= 6.2, *p* = 0.0014; interaction, F(6,78)= 1.2, *p* = 0.32. For stride time: walking speed, F(2,26)= 157.3, *p* = 3.0 × 10^− 15^; surface stiffness, F(3,39)= 4.9, *p* = 0.0053; interaction, F(6,78)= 0.85, *p* = 0.53.

### Joint-synergy recruitment showed no speed-by-stiffness interaction

CP decomposition of the joint-angle tensor extracted three rank-1 tensors, each comprising a joint synergy, an angular variation, and recruitment coefficients across participants and conditions (Figure 3). The first two joint synergies had similar structures, whereas their angular variations differed mainly in sign and peak timing (Figures 3A1, 3A2, 3B1, and 3B2). In the first synergy, the stance limb showed ankle dorsiflexion, knee extension, and hip extension after foot contact, whereas the contralateral limb showed ankle plantarflexion, knee flexion, and hip flexion; the second synergy showed the opposite pattern. Because CP decomposition does not impose orthogonality constraints, these similar but oppositely phased components may be extracted separately.

**Figure 3.**
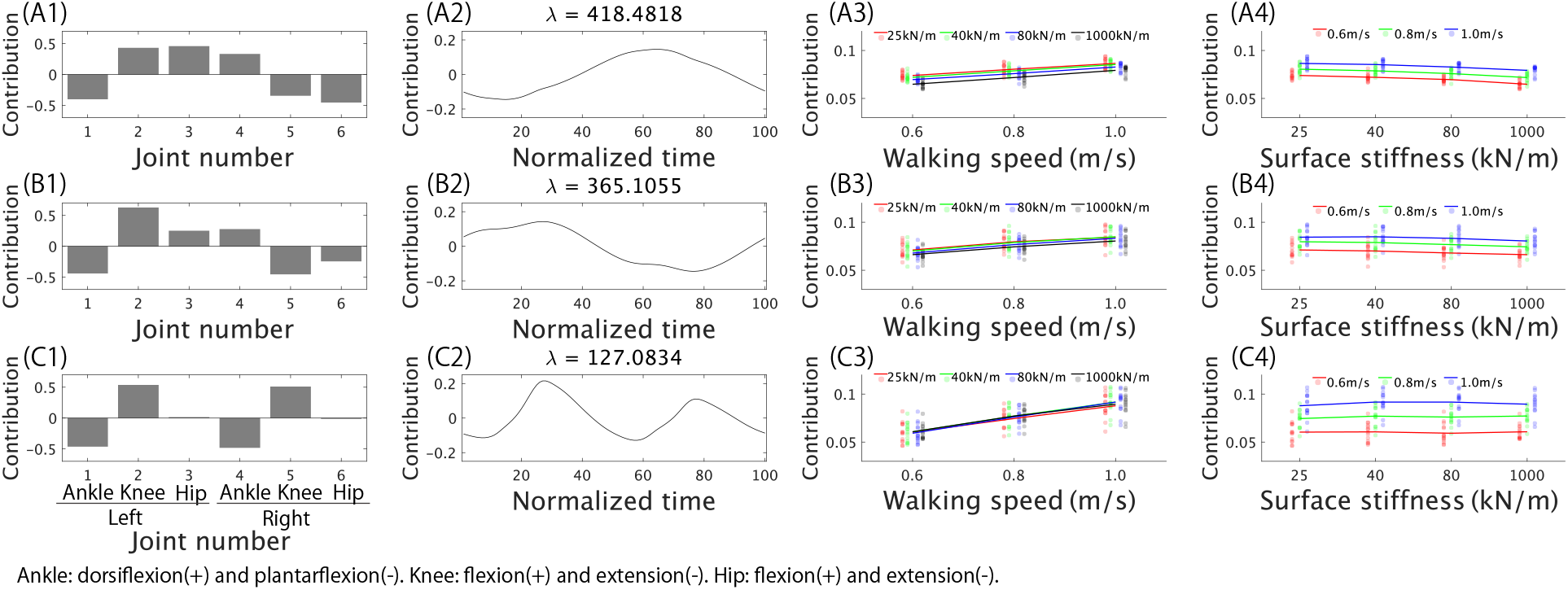
Joint-synergy recruitment shows no speed-by-stiffness interaction. (A-C) Joint rank-1 tensors extracted by CP decomposition. Within each row, panels show the joint synergy (A1, B1, and C1), corresponding angular variation (A2, B2, and C2), recruitment coefficients plotted by walking speed (A3, B3, and C3), and recruitment coefficients plotted by surface stiffness (A4, B4, and C4). In the joint-synergy panels, positive values denote ankle dorsiflexion, knee flexion, and hip flexion, whereas negative values denote ankle plantarflexion, knee extension, and hip extension. Colored lines show means across participants, and transparent points show individual participant-condition values. Two-way repeated-measures ANOVA (n = 14) showed significant main effects of walking speed for all three tensors A3: F(2, 26) = 130.6, *p* = 8.2 × 10^−14^; B3: F(2, 26) = 58.0, *p* = 7.7 × 10^−10^; C3: F(2, 26) = 119.3, *p* = 2.4 × 10^−13^), significant main effects of surface stiffness for the first two tensors (A4: F(3, 39) = 69.4, *p* = 3.2 × 10^−15^; B4: F(3, 39) = 19.4, *p* = 2.4 × 10^−7^) but not the third tensor (F(3, 39) = 2.1, *p* = 0.36), and no significant interactions (A: F(6, 78) = 1.3, *p* = 0.87; B: F(6, 78) = 0.77, *p* = 0.60 [uncorrected]; C: F(6, 78) = 2.5, *p* = 0.092). See also Figures S2 and S3.

Recruitment of these two rank-1 tensors showed significant main effects of walking speed (Figure 3A3: F(2, 26) = 130.6, p = 8.2 × 10^−14^; Figure 3B3: F(2, 26) = 58.0, p = 7.7 × 10^−10^) and surface stiffness (Figure 3A4: F(3, 39) = 69.4, p = 3.2 × 10^−15^; Figure 3B4: F(3, 39) = 19.4, p = 2.4 × 10^−7^). However, speed-by-stiffness interactions were not significant for either tensor. To aid interpretation, we merged these two joint synergies and their angular variations after weighting them by their corresponding contribution values (see Methods), and the merged component captured coordinated stance-limb extension with contralateral-limb flexion (Figures S2A and S2B).

The third joint synergy involved flexion at both ankles and extension at both knees, with an angular variation containing one positive and one negative peak (Figures 3C1 and 3C2). This angular variation was temporally aligned with the anteroposterior center-of-mass trajectory (Figures S2C-S2E), suggesting that this component may reflect forward-backward center-of-mass motion. Recruitment of this component was significantly modulated by walking speed (Figure 3C3: F(2, 26) = 119.3, p = 2.4 × 10^−13^), but not by surface stiffness (Figure 3C4: F(3, 39) = 2.1, p = 0.36), and the interaction was not significant (F(6, 78) = 2.5, p = 0.092). Thus, joint-synergy recruitment was explained by main effects of speed and, for the first two tensors, stiffness, without evidence for speed-by-stiffness interaction.

### Selected muscle-synergy recruitment showed speed-by-stiffness interactions

CP decomposition of the EMG tensor extracted five rank-1 tensors, each comprising a muscle synergy, an activity profile, and recruitment coefficients (Figure 4). The first two muscle synergies were characterized mainly by coactivation of stance-limb thigh muscles and tibialis anterior (TA), with activity profiles that were prominent from immediately after foot contact until contralateral foot contact (Figures 4A1, 4A2, 4B1, and 4B2). These profiles are consistent with muscle recruitment during weight acceptance and forward propulsion^34,35^. Recruitment of both synergies showed significant main effects of walking speed (Figure 4A3: F(2, 26) = 37.5, p = 1.1 × 10^−7^; Figure 4B3: F(2, 26) = 39.7, p = 6.3 × 10^−8^) and surface stiffness (Figure 4A4: F(3, 39) = 38.2, p = 5.4 × 10^−11^; Figure 4B4: F(3, 39) = 63.2, p = 2.4 × 10^−14^). Importantly, both synergies also showed significant speed-by-stiffness interactions (Figure 4A: F(6, 78) = 6.4, p = 1.0 × 10^−4^; Figure 4B: F(6, 78) = 7.2, p = 2.1 × 10^−5^), with stiffness-dependent differences becoming smaller at higher walking speeds.

**Figure 4.**
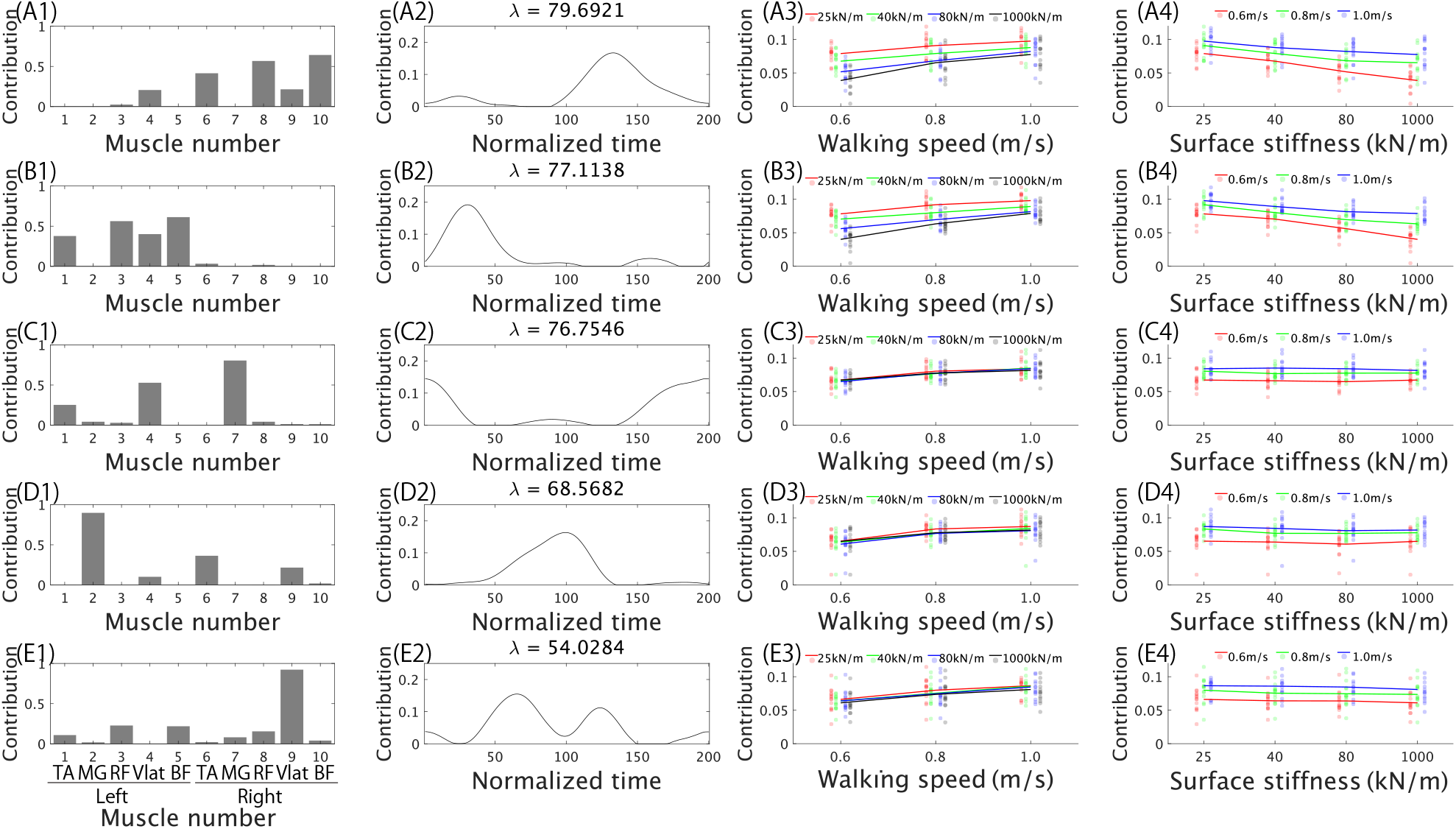
Selected muscle-synergy recruitment shows speed-by-stiffness interactions. (A-E) Muscle rank-1 tensors extracted by non-negative CP decomposition. Within each row, panels show the muscle synergy (A1, B1, C1, D1, and E1), corresponding activity profile (A2, B2, C2, D2, and E2), recruitment coefficients plotted by walking speed (A3, B3, C3, D3, and E3), and recruitment coefficients plotted by surface stiffness (A4, B4, C4, D4, and E4). Muscle labels denote tibialis anterior (TA), medial gastrocnemius (MG), rectus femoris (RF), vastus lateralis (Vlat), and biceps femoris (BF), shown for the left and right limbs. Colored lines show means across participants, and transparent points show individual participant-condition values. Two-way repeated-measures ANOVA (n = 14) showed significant main effects of walking speed for all five tensors (A3: F(2, 26) = 37.5, *p* = 1.1 × 10^−7^; B3: F(2, 26) = 39.7, *p* = 6.3 × 10^−8^; C3: F(2, 26) = 38.1, *p* = 9.3 × 10^−8^; D3: F(2, 26) = 19.7, *p* = 3.1 × 10^−5^; E3: F(2, 26) = 23.4, *p* = 7.6 × 10^−6^). Surface stiffness significantly modulated tensors A, B, and D (A4: F(3, 39) = 38.2, *p* = 5.4 × 10^−11^; B4: F(3, 39) = 63.2, *p* = 2.4 × 10^−14^; D4: F(3, 39) = 3.7, *p* = 0.0372), but not tensors C or E. Speed-by-stiffness interactions were significant for tensors A and B (A: F(6, 78) = 6.4, *p* = 1.0 × 10^−4^; B: F(6, 78) = 7.2, *p* = 2.1 × 10^−5^), but not for tensors C-E (C: F(6, 78) = 1.8, *p* = 0.53, D: F(6, 78) = 1.0, *p* = 0.41 [uncorrected], and E: F(6, 78) = 0.50, *p* = 0.81. See also Figure S4.

The third and fourth muscle synergies included medial gastrocnemius (MG) in the stance limb together with TA and vastus lateralis (Vlat) in the contralateral limb, and their activity profiles were prominent during swing and around foot contact (Figures 4C1, 4C2, 4D1, and 4D2). Recruitment of both synergies was significantly modulated by walking speed Figure 4C3: F(2, 26) = 38.1, p = 9.3 × 10^−8^; Figure 4D3: F(2, 26) = 19.7, p = 3.1 × 10^−5^). A main effect of surface stiffness was observed only for the fourth synergy (Figure 4D4: F(3, 39) = 3.7, p = 0.0372), and neither synergy showed a significant interaction. The fifth muscle synergy was dominated by right Vlat and was active around right foot contact (Figures 4E1 and 4E2). Its recruitment showed a significant main effect of speed (Figure 4E3: F(2, 26) = 23.4, p = 7.6 × 10^− 6^), but neither the main effect of surface stiffness nor the interaction was significant (Figure 4E). Thus, the EMG data contained both independently modulated components (Figures 4C-4E) and interactively modulated components (Figures 4A and 4B).

### Subject-wise and rank-sensitivity analyses confirmed the main components

We next examined whether the rank-1 tensors were stable when CP decomposition was applied separately to each participant. For joint-angle data, all three joint synergies, angular variations, and recruitment patterns extracted from individual participants were consistent with those extracted from the pooled dataset (Figure S3). For EMG data, four of the five rank-1 tensors showed consistent muscle synergies, activity profiles, and recruitment patterns between subject-wise and pooled analyses (Figure S4). The fifth EMG tensor was less stable: its muscle synergy and activity profile were not consistent across analyses, although its recruitment coefficients were correlated between analyses.

We also tested the sensitivity of the results to the number of rank-1 tensors (R; Figure S5). For joint-angle data, tensors similar to those in Figure 3 were recovered when R was varied, indicating that the three joint components were stable across rank choices. For EMG data, tensors similar to those in Figures 4A-4D appeared when fewer components were extracted and were also recovered at higher ranks, whereas the fifth tensor was less robust. These analyses indicate that the three joint tensors and four of the five EMG tensors represent the major components of the data.

### Conventional matrix factorization reproduced the joint-versus-muscle contrast

Finally, we compared CP decomposition with conventional matrix factorization approaches (Figure 5). When PCA and NNMF were applied separately to each participant and condition, these methods extracted joint or muscle synergies and their temporal profiles, but they did not directly provide condition-dependent recruitment coefficients (Figures 5A and 5B). When the data were instead arranged as matrices with dimensions (joint or muscle × time) × condition, PCA and NNMF extracted spatiotemporal synergies and their recruitment across conditions (Figures 5C and 5D). The recruitment of joint spatiotemporal synergies showed significant main effects of speed (F(2, 26) = 198.0, p = 1.8 × 10^−16^) and surface stiffness (F(3, 39) = 38.8, p = 8.7 × 10^−12^), but no interaction (F(6, 78) = 0.58, p = 0.74; Figure 5C). In contrast, the recruitment of muscle spatiotemporal synergies showed significant main effects of speed (F(2, 26) = 128.5, p = 3.3 × 10^−14^) and surface stiffness (F(3, 39) = 41.3, p = 3.4 × 10^−12^), as well as a significant interaction (F(6, 78) = 5.5, p = 8.8 × 10^−5^; Figure 5D). These results are consistent with the CP-decomposition findings: walking speed and surface stiffness affected joint-level coordination primarily through independent main effects, whereas selected muscle-level components showed interactive modulation.

**Figure 5.**
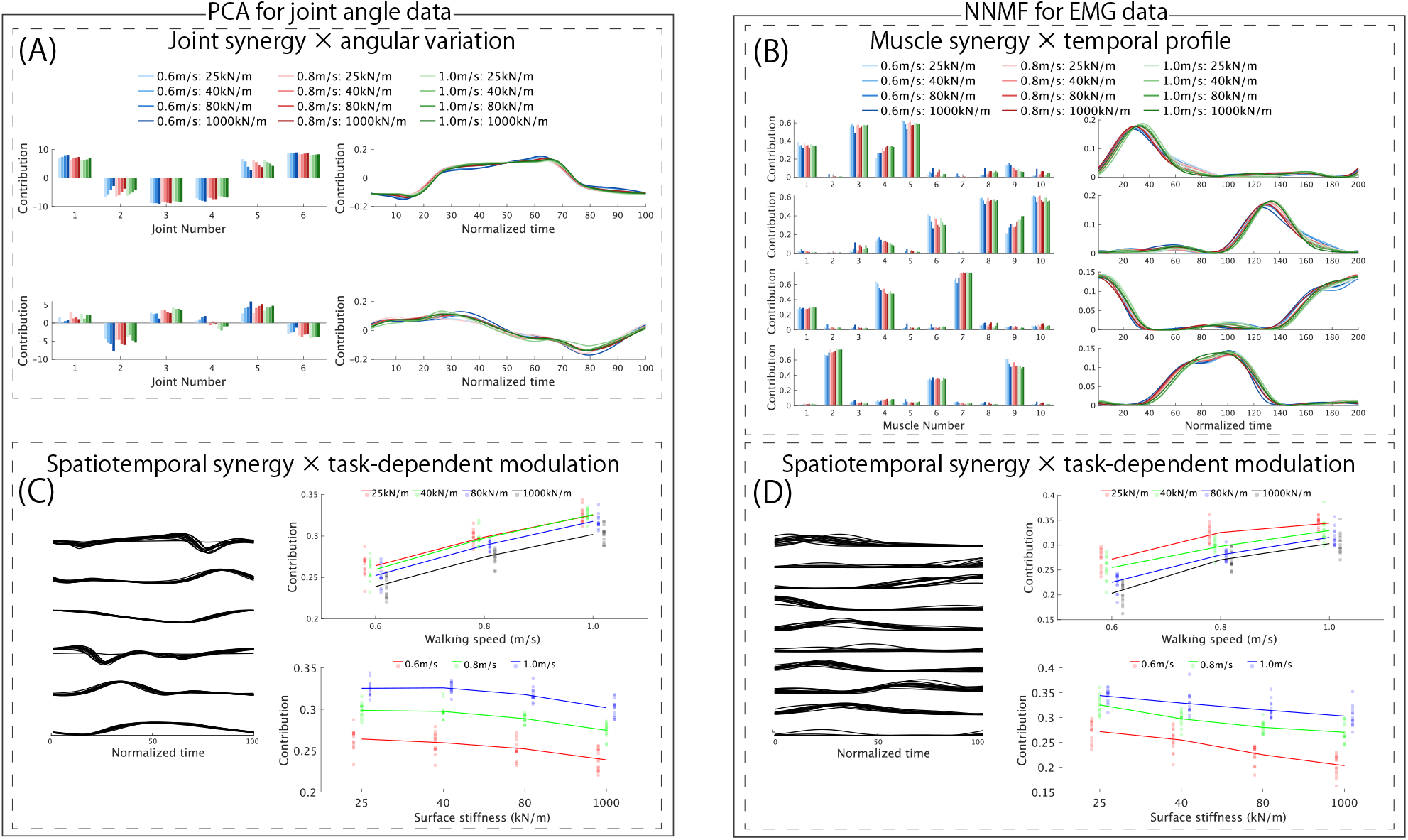
Conventional matrix factorization reproduces the joint-versus-muscle contrast. (A and B) PCA applied to joint-angle data (A) and NNMF applied to EMG data (B) separately for each participant and condition. Left panels show spatial synergies, and right panels show temporal components. (C and D) PCA or NNMF applied to matrices arranged as (joint or muscle x time) x condition within each participant. Left panels show spatiotemporal synergies, and right panels show condition-dependent recruitment coefficients. Colored lines show means across participants, and transparent points show individual participant-condition values. For joint spatiotemporal synergies (C), two-way repeated-measures ANOVA (n = 14) showed significant main effects of walking speed ((F(2, 26) = 198.0, *p* = 1.8 × 10^−16^) and surface stiffness (F(3, 39) = 38.8, *p* = 8.7 × 10^−12^), with no significant interaction (F(6, 78) = 0.58, *p* = 0.74). For muscle spatiotemporal synergies (D), the main effects of walking speed (F(2, 26) = 128.5, *p* = 3.3 × 10^−14^) and surface stiffness (F(3, 39) = 41.3, *p* = 3.4 × 10^−12^), as well as their interaction (F(6, 78) = 5.5, *p* = 8.8 × 10^−5^), were significant.

## DISCUSSION

Using an open dataset of controlled-speed walking on a variable-stiffness treadmill^33^, we examined whether gait speed and surface stiffness modulate gait independently or interactively. Stride length, stride time, and joint-synergy recruitment were primarily shaped by independent main effects of speed and stiffness, whereas two of the five muscle synergies showed speed-by-stiffness interactions. Previous studies have reported that stride parameters vary with walking speed^36^ and with surface stiffness when speed is not controlled^37^. Speed-dependent modulation has also been reported for both joint synergies^3,18^ and muscle synergies^10,18^. The present study extends these findings by showing that, under controlled-speed conditions, surface stiffness influences stride and joint-level coordination without interacting with gait speed, whereas it can interactively modulate selected muscle-synergy recruitment patterns. Because walking speed can confound gait comparisons^24–26^, the controlled design of the dataset combined with CP decomposition was critical for separating speed-dependent from stiffness-dependent effects.

The interaction observed for the first two muscle synergies may have implications for walking training and rehabilitation. These synergies were associated with weight acceptance and forward propulsion^34,35^, and their recruitment was greater on softer surfaces at slower walking speeds, whereas stiffness-dependent differences became smaller at higher walking speeds (Figures 4A and 4B). These findings raise the possibility that softer surfaces could be used to increase selected muscle-synergy recruitment at slower walking speeds, although this interpretation requires direct testing in clinical populations. As walking speed increases, however, the additional effect of surface compliance may become less pronounced. This interpretation should be tested directly in clinical populations because the present dataset included only healthy young adults.

For the step-related joint synergies (Figures 3A, 3B, S2A, and S2B), recruitment increased as the surface became softer. This pattern may reflect greater joint excursions and is consistent with increased hip, knee, and ankle ranges of motion on compliant surfaces^22^. It may also correspond to a proactive response to reduce tripping risk on softer surfaces, such as dorsiflexing the ankle and increasing toe clearance^19,20^. The present results suggest that this kinematic strategy depends on surface stiffness but not on its interaction with walking speed.

By contrast, the third joint synergy (Figure 3C) appeared to reflect anteroposterior CoM motion rather than stiffness-related adaptation. Its angular variation was temporally aligned with the anteroposterior CoM trajectory (Figures S2C–S2E), and its recruitment was modulated by walking speed but not by surface stiffness or the speed-by-stiffness interaction. This pattern suggests that anteroposterior CoM-related joint coordination is largely governed by walking speed under the tested conditions.

At the muscle-synergy level, the stiffness-dependent recruitment of the first two muscle synergies is consistent with previous reports of greater muscle activity on compliant surfaces^19,20,22^. The third and fourth muscle synergies were structurally left-right symmetric, but only the fourth showed a main effect of surface stiffness. This asymmetry in statistical modulation may be related to previous observations that, when one leg walks on a soft surface and the other on a hard surface, some muscles do not necessarily show significant left-right differences^19^. In addition, the fifth rank-1 tensor should be interpreted cautiously because its muscle synergy and activity profile were not stable between pooled-subject and subject-wise analyses.

Methodologically, CP decomposition was useful for this question because it separated spatial synergies, temporal components, and condition-dependent recruitment within a single framework. Matrix factorization methods such as PCA and NNMF can identify spatial and temporal components but do not directly estimate condition-dependent recruitment when applied separately to each condition. More flexible tensor approaches, including Tucker decomposition^29^ and matrix tri-factorization^38,39^, may be advantageous in some situations; however, they require multiple rank parameters, and parameter combinations that explain a similar amount of variance may not be unique^32^. CP decomposition is less flexible but requires specification of only a single rank parameter, making it a practical extension of PCA and NNMF for comparing task-dependent synergy modulation.

### Limitations of the study

First, the analyzed dataset included EMG recordings from only 10 lower-limb muscles, which prevented evaluation of muscles commonly assessed in gait studies, such as the soleus, gluteus maximus, and trunk muscles^5,18^. Previous studies have reported increased activity of the soleus^19,20,22^ and erector spinae^20^ during compliant-surface walking. Therefore, the present results do not determine whether gait speed and surface stiffness affect these unrecorded muscles independently or interactively.

Second, the open dataset was limited to young healthy adults, and the final analysis included seven female and seven male participants after EMG-based exclusions. The dataset therefore does not allow strong conclusions about aging, clinical populations, or sex differences. Replication in larger and more diverse cohorts will be needed, especially because comparable controlled-speed, variable-stiffness datasets are currently scarce.

Third, we focused on steady-state gait by analyzing the last 30 qualifying gait cycles in each condition. This design was appropriate for separating speed and stiffness effects during stable walking but did not address responses to abrupt changes in surface stiffness. Previous studies have examined transitions such as stepping onto a compliant surface with one leg^20,23^ or taking only a few steps on a compliant surface with both legs^19,21^. Future analyses of early gait cycles in the present dataset could complement the steady-state analysis by characterizing transient responses, although such responses may also include stumbling, startle effects, or left-right asymmetry.

Finally, because the source protocol used a fixed progression of speed trials and a fixed order of stiffness conditions within each speed trial, residual order, fatigue, or adaptation effects cannot be fully excluded.

However, the analysis focused on the last 30 qualifying gait cycles in each condition to characterize steady-state behavior.

## RESOURCE AVAILABILITY

### Lead contact

Requests for further information and resources should be directed to and will be fulfilled by the lead contact, Ken Takiyama.

### Materials availability

This study did not generate new unique reagents.

### Data and code availability

- The source dataset analyzed in this study is publicly available in Figshare at doi:10.6084/m9.figshare.27180288.
- The MATLAB scripts used to preprocess the data, perform the analyses, and generate the figures will be deposited in Figshare and made publicly available as of the date of publication. During peer review, the scripts are available from the lead contact upon reasonable request.

## ACKNOWLEDGMENTS

This work was supported by a Grant-in-Aid for Scientific Research (B) to KT (24K02840).

## AUTHOR CONTRIBUTIONS

Conceptualization, T.I. and K.T.; methodology, T.I. and K.T.; investigation, T.I. and K.T.; writing—original draft, T.I.; writing—review & editing, K.T.; funding acquisition, K.T.; resources, T.I. and K.T.; supervision, K.T. All authors have read and approved the final version of the manuscript.

## DECLARATION OF GENERATIVE AI AND AI-ASSISTED TECHNOLOGIES

During the preparation of this work, the authors used ChatGPT (OpenAI) to improve language quality and readability. After using this tool, the authors reviewed and edited the content as needed and take full responsibility for the content of the publication.

## SUPPLEMENTAL INFORMATION

**Document S1 and Figures S1–S5**.

## STAR★METHODS

### KEY RESOURCES TABLE

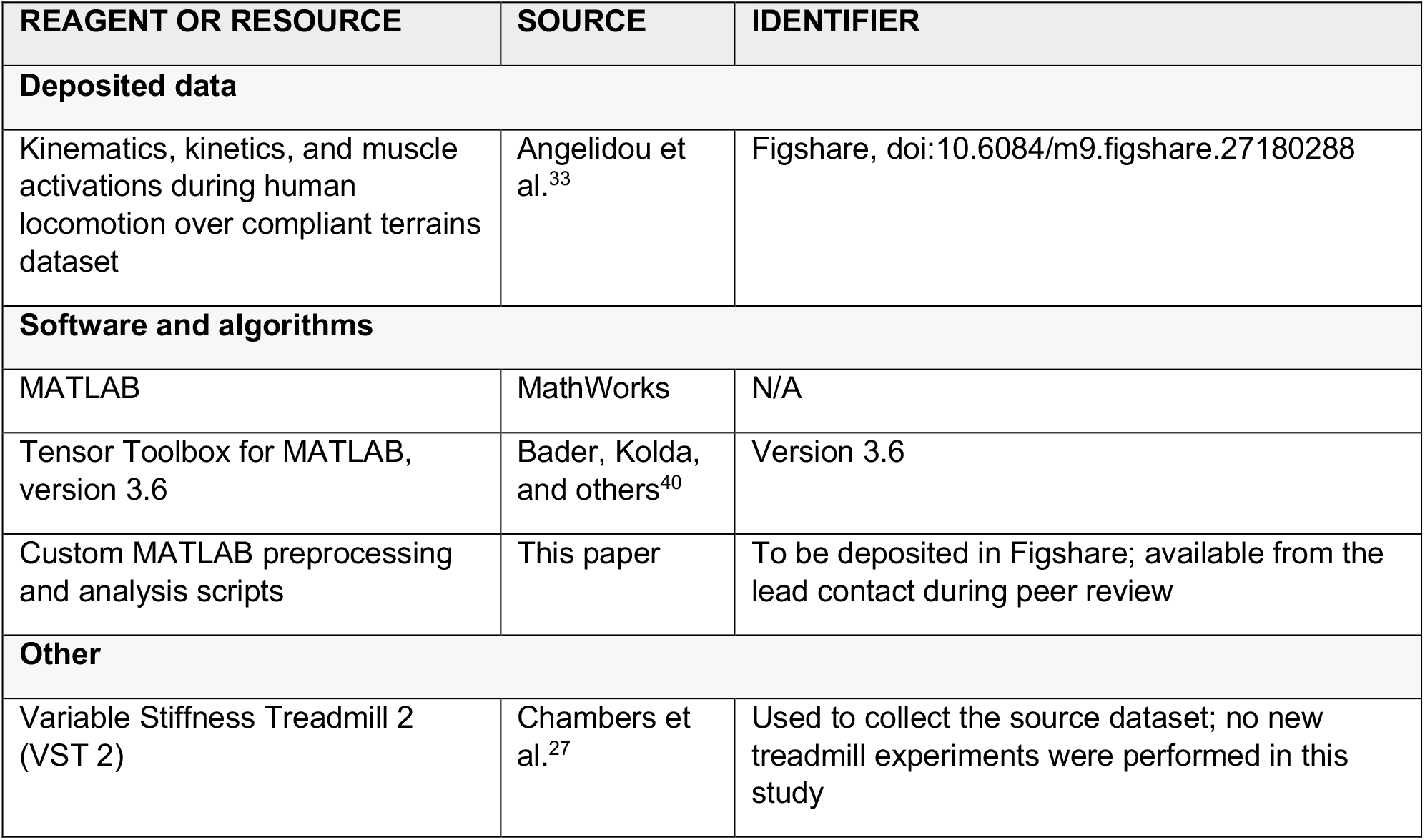

| REAGENT OR RESOURCE | SOURCE | IDENTIFIER |
| --- | --- | --- |
| **Deposited data** |  |  |
| Kinematics, kinetics, and muscle activations during human locomotion over compliant terrains dataset | Angelidou et al.<sup>33</sup> | Figshare, doi:10.6084/m9.figshare.27180288 |
| **Software and algorithms** |  |  |
| MATLAB | MathWorks | N/A |
| Tensor Toolbox for MATLAB, version 3.6 | Bader, Kolda, and others<sup>40</sup> | Version 3.6 |
| Custom MATLAB preprocessing and analysis scripts | This paper | To be deposited in Figshare; available from the lead contact during peer review |
| **Other** |  |  |
| Variable Stiffness Treadmill 2 (VST 2) | Chambers et al.<sup>27</sup> | Used to collect the source dataset; no new treadmill experiments were performed in this study |

## EXPERIMENTAL MODEL AND STUDY PARTICIPANT DETAILS

### Human participants in the source dataset

This study reanalyzed an existing open human gait dataset collected during walking on a variable-stiffness treadmill^33^. The source dataset included 20 healthy adults (10 female and 10 male; age range, 19-37 years), with demographic and anthropometric information reported in the original publication. No new human-participant data were collected in the present study. After the EMG quality-control procedures described below, six participants were excluded, and the final analysis included 14 participants (7 female and 7 male; age range, 19-37 years). Ethics approval and informed consent procedures for the source dataset are described in Angelidou et al.^33^. No new human-participant data were collected in the present study.

## METHOD DETAILS

### Experimental protocol and analyzed gait cycles

The source dataset contained electromyography (EMG) and joint-angle data recorded during treadmill walking under 12 combinations of walking speed and surface stiffness. The walking speeds were 0.6, 0.8, and 1.0 m/s, and the surface-stiffness levels were 25, 40, 80, and 1000 kN/m. For each speed trial, participants first completed a rigid warm-up condition and then walked at the four stiffness levels; each condition contained 150 gait cycles, with 5-min rests between speed trials. We analyzed steady-state gait by selecting the last 30 gait cycles that satisfied the inclusion criteria from the latter 100 gait cycles in each condition. A gait cycle was defined as the interval from left heel strike to the subsequent left heel strike. Stride length, stride time, joint-angle trajectories, and EMG profiles were averaged across the selected 30 gait cycles for each participant and condition.

### Preprocessing of joint-angle data

Three-dimensional marker positions were recorded at 100 Hz in the source dataset, and joint angles were calculated using Vicon’s Plug-in Gait Full-body model^33^. We analyzed six sagittal-plane joint angles: left and right ankle, knee, and hip angles. Each gait cycle was time-normalized to 100 samples. For each participant and condition, joint-angle trajectories were averaged across the selected 30 gait cycles. For CP decomposition, each joint-angle trajectory was standardized within each participant and joint across all conditions so that the mean and standard deviation were 0 and 1, respectively. The resulting joint-angle tensor had dimensions 6 x 100 x 168, where 168 corresponds to 14 participants x 12 conditions.

### Preprocessing of EMG data

Surface EMG signals were recorded at 2000 Hz bilaterally from 10 lower-limb muscles: tibialis anterior (TA), medial gastrocnemius (MG), rectus femoris (RF), vastus lateralis (Vlat), and biceps femoris (BF)^41^. EMG processing followed the custom MATLAB script provided with the source dataset, with an additional demeaning step. Raw EMG signals were band-pass filtered at 30-300 Hz using a fourth-order zero-phase Butterworth filter, demeaned, and full-wave rectified. The signal envelope was then estimated using a moving root-mean-square window of 200 samples and low-pass filtered at 5 Hz using a fourth-order zero-phase Butterworth filter. We did not use the MVC-based normalization reported in the source dataset; instead, after time normalization and averaging, EMG data were min-max normalized within each participant and muscle across all conditions.

### EMG quality control and participant exclusion

Each EMG gait cycle was time-normalized to 200 samples. To exclude noisy EMG profiles, each gait cycle in the latter 100 gait cycles of each condition was evaluated using four criteria: 95th percentile of EMG amplitude, root-mean-square amplitude, timing of the peak EMG amplitude, and cosine similarity to a template EMG profile. The template for each muscle and condition was defined as the median EMG profile across gait cycles in the latter half of that condition. The 95th-percentile and root-mean-square criteria were used to detect spike-like noise and sustained high-amplitude noise, respectively; peak timing was used to detect channel swaps or isolated spikes; and cosine similarity was used to detect profiles that deviated from the typical gait-phase pattern^42,40^. Outliers were identified using MATLAB’s isoutlier function with the median option for each criterion. A gait cycle was excluded when it was identified as an outlier by more than one criterion.

Participants were excluded in two stages. First, one participant (subject #15) had fewer than 30 analyzable gait cycles and was excluded. Three additional participants (subjects #2, #11, and #20) were excluded because their numbers of excluded gait cycles were more than one standard deviation above the across-participant mean. Second, because the above criteria were applied within participants and may not detect globally noisy EMG profiles, we calculated between-participant correlation coefficients of post-processed EMG profiles averaged across the selected 30 gait cycles for each condition. For each participant, the lowest correlation coefficient across muscles was quantified and then averaged across conditions; participants whose averaged value was more than one standard deviation below the across-participant mean were excluded (subjects #3 and #8). The resulting EMG tensor had dimensions 10 x 200 x 168. Excluded EMG profiles are shown in Figure S1.

### Tensor construction and CP decomposition

CP decomposition was applied separately to the joint-angle tensor and the EMG tensor. For the joint-angle tensor ***X***, the k-th participant-condition slice was approximated as follows:

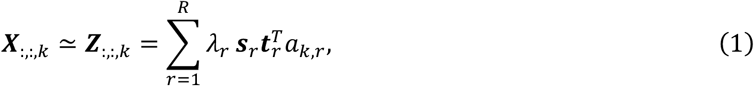

where ***s***_*r*_ = (*s*_1,*r*_, *s*_2,*r*_, …, *s*_6,*r*_) is the *r*-th joint synergy, ***t***_*r*_ = (*t*_1,*r*_, *t*_2,*r*_, …, *t*_100,*r*_) is the corresponding angular variation, *a*_*k,r*_ is the recruitment coefficient for participant-condition, *k, λ*_*r*_ ≥ 0 is the contribution of the *r*-th rank-1 tensor, and *k* = 1, …, 168 indexes participant-condition pairs. The factor vectors were normalized to unit norm, and repeated-measures analyses were applied to the recruitment coefficients after sorting them by participant and condition.

For the non-negative EMG tensor ***Y***, the *k*-th participant-condition slice was approximated as follows:

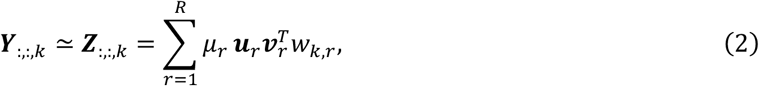

where ***u***_*r*_ = (*u*_1,*r*_, *u*_2,*r*_, …, *u*_10,*r*_) is the *r*-th muscle synergy, ***v***_*r*_ = (*v*_1,*r*_, *v*_2,*r*_, …, *v*_200,*r*_) is the corresponding activity profile, *w*_*k,r*_ is the recruitment coefficient, and all elements of ***u***_*r*_, ***v***_*r*_, and the recruitment factor *w*_*k,r*_ were constrained to be non-negative. Joint-angle decomposition was performed using cp_als, and EMG decomposition was performed using cp_nmu; both functions were implemented in Tensor Toolbox for MATLAB^18,26,32^.

Each decomposition was repeated with 100 random initializations, and the solution with the lowest reconstruction error was selected. The convergence tolerance and maximum number of iterations were set to 10^™6^ and 100, respectively.

### Determination of the number of rank-1 tensors

The number of rank-1 tensors R was determined as the minimum number of components that explained more than 70% of the variance of the original data. Variance explained was defined as Var(***Z***)/Var(***X***) for the joint-angle tensor and Var(***Z***)/Var(***Y***) for the EMG tensor, where ***Z*** denotes the tensor reconstructed from the extracted rank-1 tensors. This metric was used to apply a consistent criterion to joint-angle and EMG data, following previous CP-decomposition studies^26,32^. It differs from the conventional variance accounted for (VAF) often used in NNMF; in the present EMG analysis, the 70% criterion corresponded to approximately 86% in conventional VAF terms.

### Merged joint synergy shown in Figure S2

For interpretive visualization of the first two joint rank-1 tensors, we calculated the merged joint synergy and merged angular variation shown in Figures S2A and S2B. The joint synergy and angular variation of each rank-1 tensor were weighted by 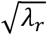, and the weighted components were then summed across the two tensors. This allocation was used because the contribution coefficient in CP decomposition weights the product of the spatial, temporal, and recruitment factors, making its assignment to individual factors non-unique.

### Similarity index and subject-wise CP decomposition

To compare rank-1 tensors extracted from the pooled data with those extracted from each participant separately, we decomposed each participant’s data using the same rank as in the pooled analysis (*R* = 3 for joint-angle data and *R* = 5 for EMG data). Components were matched using a similarity index defined as the average of the absolute correlation coefficients for the spatial synergy, temporal component, and recruitment coefficients. A similarity index greater than 0.75 was interpreted as indicating consistent components, following previous studies^26,32^.

For joint-angle data, where signs are arbitrary, rank-1 tensors were reordered according to the similarity index and their signs were adjusted. If the correlation of recruitment coefficients was negative, the recruitment and joint-synergy factors were multiplied by -1. If the correlations for the joint synergy and angular variation were both negative, the joint-synergy and angular-variation factors were multiplied by -1. For EMG data, signs were not adjusted because all factors were constrained to be non-negative. Rank sensitivity was also assessed by repeating CP decomposition with alternative values of *R* and calculating similarity indices between the selected components and the components extracted at other ranks (Figure S5).

### Comparison with PCA and NNMF

For comparison with conventional matrix factorization, PCA was applied to joint-angle data and NNMF was applied to EMG data in two ways. First, PCA or NNMF was applied separately to each participant and condition using matrices with dimensions corresponding to joints or muscles × normalized time. For this analysis, joint-angle data were standardized within each participant and condition so that each joint had a mean of 0 and a standard deviation of 1, whereas EMG data were min-max normalized within each participant and condition.

Second, PCA or NNMF was applied to matrices arranged as (joint or muscle × time) × condition for each participant. In this analysis, PCA extracted joint spatiotemporal synergies and NNMF extracted muscle spatiotemporal synergies together with condition-dependent recruitment coefficients. The number of components was chosen using the same variance-explained criterion as that used for CP decomposition.

## QUANTIFICATION AND STATISTICAL ANALYSIS

Statistical analyses were performed on data from 14 participants. Repeated-measures ANOVA (rANOVA) was applied to stride length, stride time, the recruitment coefficients of each joint and muscle rank-1 tensor, and the recruitment coefficients obtained from PCA/NNMF-based spatiotemporal analyses. Walking speed (0.6, 0.8, and 1.0 m/s) and surface stiffness (25, 40, 80, and 1000 kN/m) were treated as within-subject factors. Paired t-tests were used to test whether correlations between components extracted from pooled and subject-wise decompositions were greater than zero. Significance was defined as an adjusted p value < 0.05. P values were corrected using Bonferroni correction unless otherwise specified. Exact F statistics, degrees of freedom, and p values are reported in the Results and figure legends.

## ADDITIONAL RESOURCES

No additional resources are reported in this study.

