## Supplementary material for "Gait speed and surface stiffness interactively modulate selected muscle- but not joint-synergy recruitment during compliant-surface walking": DocumentS1

### EMG data quality control

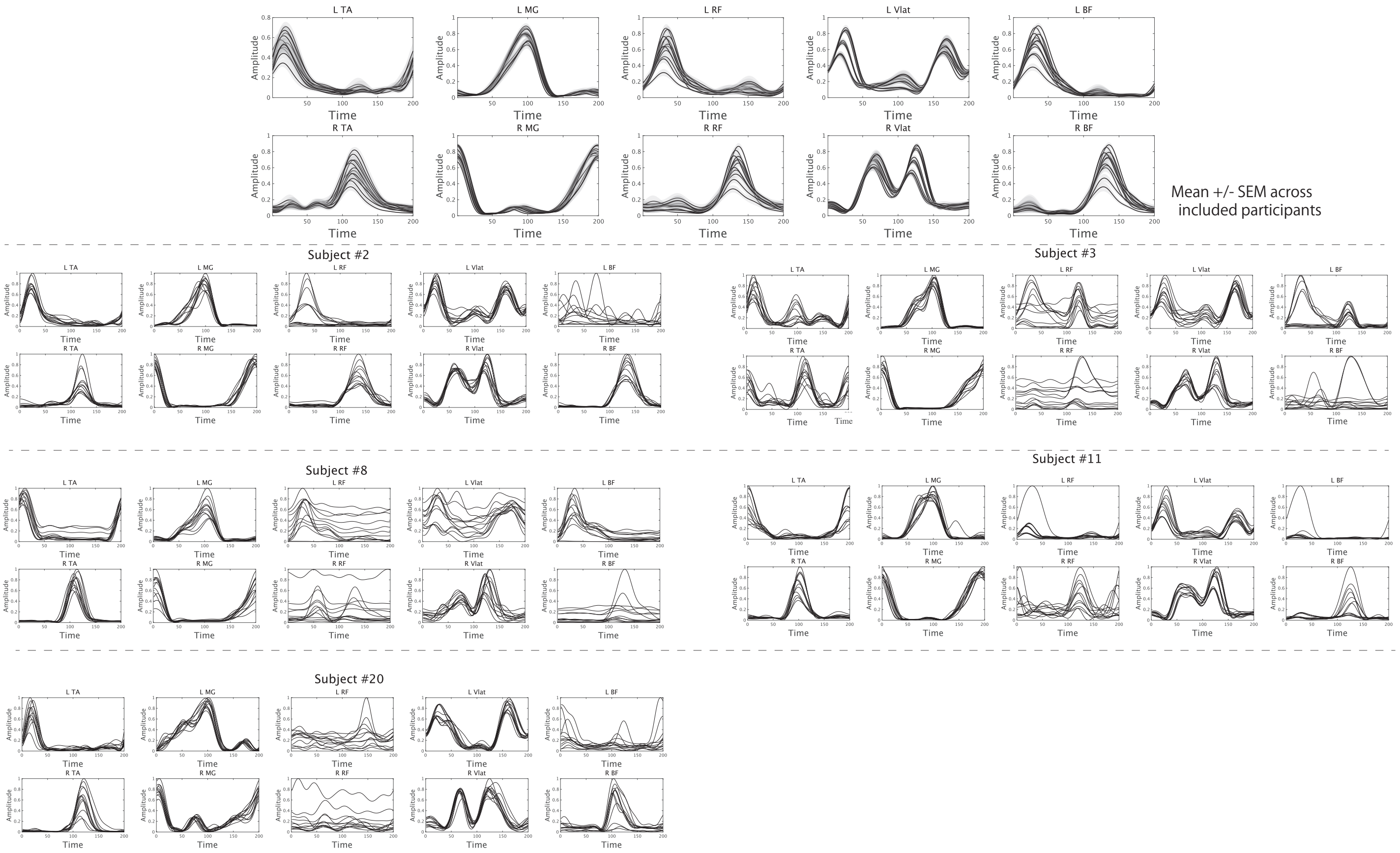


Figure S1. EMG quality-control criteria and excluded EMG profiles

Top panels show mean EMG profiles (solid lines) and SEM (shaded areas) across included participants after exclusions. Lower panels show excluded EMG profiles. Subject #2 was excluded because the number of outlier gait cycles exceeded one standard deviation above the across-participant mean, with noisy profiles particularly evident in left BF. Subjects #3 and #8 were excluded because the between-participant correlations of EMG profiles were more than one standard deviation below the across-participant mean, with noisy profiles particularly evident in left RF, right RF, and right BF for subject #3, and in right RF and right BF for subject #8. Subjects #11 and #20 were excluded because the number of outlier gait cycles exceeded one standard deviation above the across-participant mean; noisy profiles were particularly evident in right RF and high-amplitude profiles in left RF and left BF for subject #11, and in left RF, left BF, and right RF for subject #20. Subject #15 is not shown because fewer than 30 analyzable gait cycles were available.

**Merged joint synergy and angular variation**

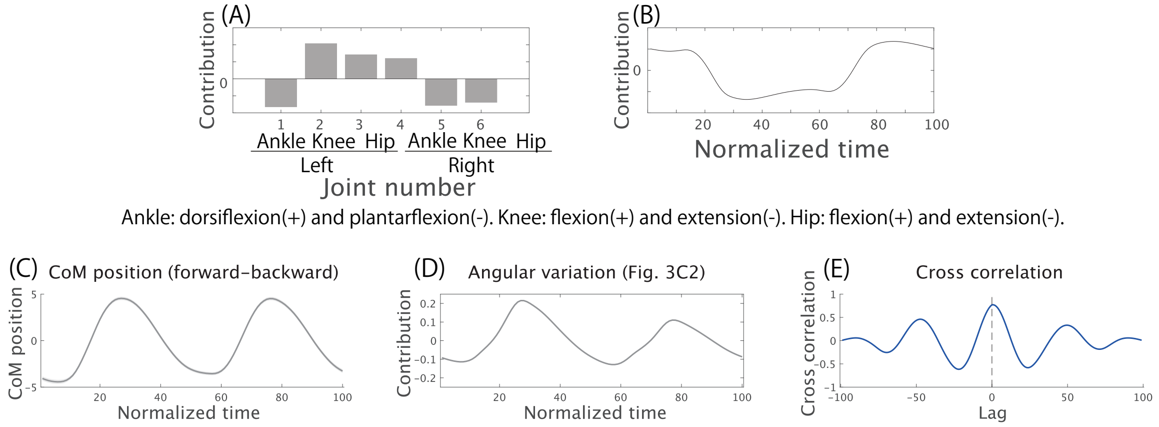


Figure S2. Merged joint synergy and relationship between angular variation and center-of-mass position

(A) Merged joint synergy calculated from the rank-1 tensors shown in Figures 3A and 3B.

(B) Merged angular variation calculated from the rank-1 tensors shown in Figures 3A and 3B.

(C) Anteroposterior center-of-mass (CoM) position. The solid line indicates the mean across participants and conditions, and the shaded area indicates SEM.

(D) Angular variation shown in Figure 3C2.

(E) Cross-correlation between the anteroposterior CoM position and the angular variation shown in Figure 3C2. The solid line indicates the mean across participants and conditions, the shaded area indicates SEM, and the vertical dotted line indicates zero lag.

### Subject-wise CP decomposition of joint-angle data

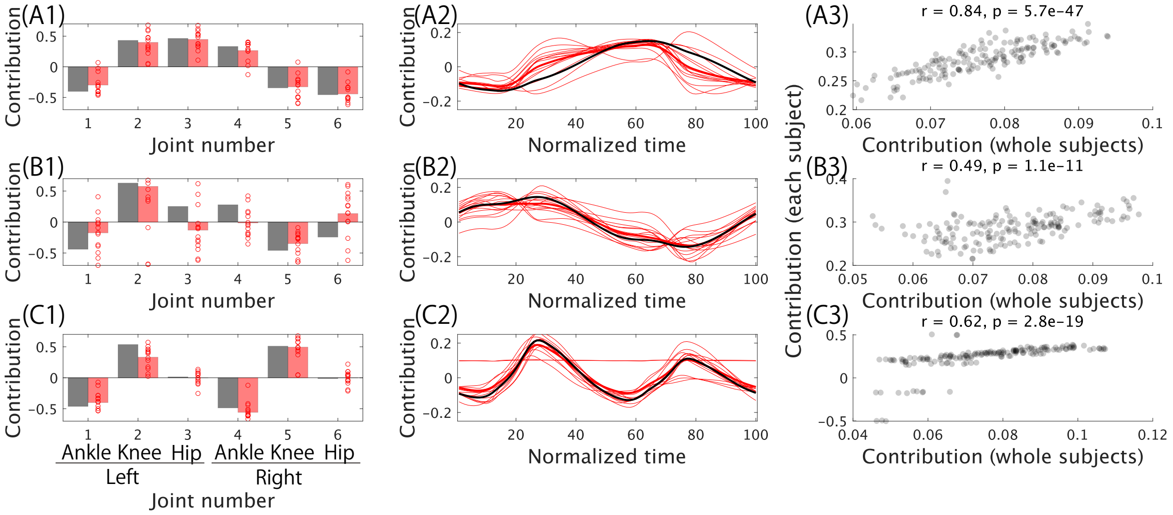


Ankle: dorsiflexion(+) and plantarflexion(-). Knee: flexion(+) and extension(-). Hip: flexion(+) and extension(-).

Figure S3. Subject-wise validation of rank-1 tensors extracted from joint-angle data

Joint synergies, angular variations, and recruitment coefficients extracted from each participant are compared with those extracted from the pooled dataset. Components shown in the same row correspond to the same rank-1 tensor. (A1, B1, and C1) Red circles and bars indicate the joint synergies for individual participants and their across-participant mean, respectively; black bars indicate the joint synergies shown in Figure 3. (A2, B2, and C2) Thin and thick red lines indicate the angular variations for individual participants and their across-participant mean, respectively; black lines indicate the angular variations shown in Figure 3. (A3, B3, and C3) Correlations between recruitment coefficients estimated from each participant separately and those estimated from the pooled dataset. The correlations were significant in A3 (r = 0.84, $p = 5.7 \times 10^{-47}$), B3 (r = 0.49, $p = 1.1 \times 10^{-11}$), and C3 (r = 0.62, $p = 2.8 \times 10^{-19}$).

**Subject-wise CP decomposition of EMG data**

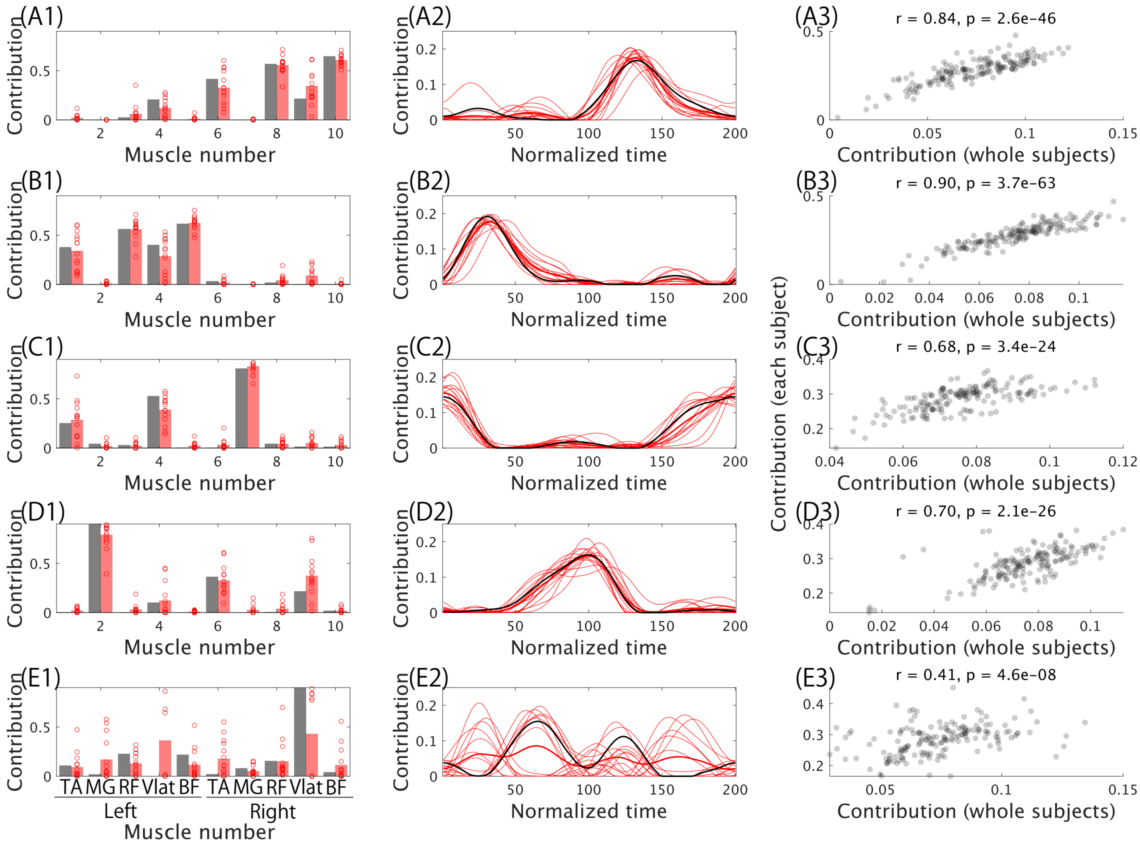


Figure S4. Subject-wise validation of rank-1 tensors extracted from EMG data

Muscle synergies, activity profiles, and recruitment coefficients extracted from each participant are compared with those extracted from the pooled dataset. Components shown in the same row correspond to the same rank-1 tensor. (A1, B1, C1, D1, and E1) Red circles and bars indicate the muscle synergies for individual participants and their across-participant mean, respectively; black bars indicate the muscle synergies shown in Figure 4. (A2, B2, C2, D2, and E2) Thin and thick red lines indicate the activity profiles for individual participants and their across-participant mean, respectively; black lines indicate the activity profiles shown in Figure 4. (A3, B3, C3, D3, and E3) Correlations between recruitment coefficients estimated from each participant separately and those estimated from the pooled dataset. The correlations were significant in A3 (r = 0.84, $p = 2.6 \times 10^{-46}$), B3 (r = 0.90, $p = 3.7 \times 10^{-63}$), C3 (r = 0.68, $p = 3.4 \times 10^{-24}$), D3 (r = 0.70, $p = 2.1 \times 10^{-26}$), and E3 (r = 0.41, $p = 4.6 \times 10^{-8}$).

### Rank-sensitivity analysis

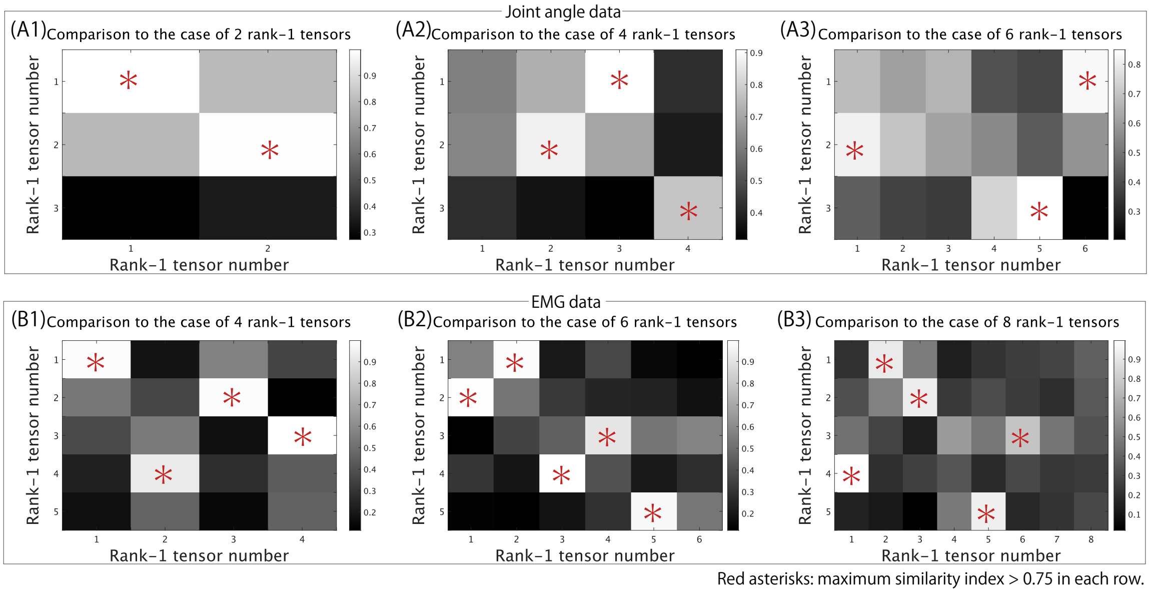


Figure S5. Similarity indices across different numbers of rank-1 tensors

(A1-A3) Similarity indices for joint-angle rank-1 tensors compared with decompositions using $R = 2$ (A1), $R = 4$ (A2), and $R = 6$ (A3).

(B1-B3) Similarity indices for EMG rank-1 tensors compared with decompositions using $R = 4$ (B1), $R = 6$ (B2), and $R = 8$ (B3). Red asterisks indicate pairs with a similarity index greater than 0.75 and the maximum similarity index within each row.
